# Site-directed mutagenesis of the 1,3-β glucan synthase catalytic subunit of *Pneumocystis jirovecii* and susceptibility assays suggest its sensitivity to caspofungin

**DOI:** 10.1101/340281

**Authors:** A. Luraschi, S. Richard, P. M. Hauser

## Abstract

The echinocandin caspofungin inhibits the catalytic subunit Gsc1 of the enzymatic complex synthetizing 1,3-β glucan, an essential compound of the fungal wall. Studies in rodents showed that caspofungin can treat *Pneumocystis* infections. However, its efficacy against *Pneumocystis jirovecii*, the species infecting exclusively humans, remains controversial. The aim of this study was to assess the sensitivity to caspofungin of the *P. jirovecii* Gsc1 subunit, as well as of those of *Pneumocystis carinii* and *Pneumocystis murina* infecting respectively rats and mice. In absence of an established *in vitro* culture method for *Pneumocystis* species, we used functional complementation of the *Saccharomyces cerevisiae gsc1* deletant. In the fungal pathogen *Candida albicans*, mutations leading to amino acid substitutions in Gsc1 confer resistance to caspofungin. We introduced the corresponding mutations into the *Pneumocystis gsc1* genes using site-directed mutagenesis. In spot dilution tests, the sensitivity to caspofungin of the complemented strains decreased with the number of mutations introduced, suggesting that the wild-type enzymes are sensitive. The minimum inhibitory concentrations of caspofungin determined by E-test^®^ and Yeastone^®^ for strains complemented with *Pneumocystis* enzymes (respectively 0.125 and 0.12 microg/ml) were identical to those upon complementation with the enzyme of *C. albicans* that is sensitive to caspofungin. However, they were lower than the MICs upon complementation with the enzyme of the resistant species *Candida parapsilosis* (0.19 and 0.25). Sensitivity levels of Gsc1 enzymes of the three *Pneumocystis* species were similar. Our results suggest that *P. jirovecii* is sensitive to caspofungin during infections, as *P. carinii* and *P. murina*.

## INTRODUCTION

The *Pneumocystis* genus comprises fungal species that colonize the lungs of mammals (1–4). Each of them displays strict host specificity for a single mammalian species. The species infecting humans is *Pneumocystis jirovecii*, an opportunistic pathogen that can cause fatal pneumonia if not treated (*<u>P</u>neumocystis* <u>p</u>neumonia, PCP). The most efficient drug against *P. jirovecii* is currently cotrimoxazole, a combination of sulfamethoxazole and trimethoprim, two inhibitors of enzymes that are involved in the folic acid biosynthesis pathway. However, resistance is emerging due to the selection of isolates carrying specific mutations in the active site of the targets of both molecules (5–9). Moreover, cotrimoxazole can cause important side effects in some patients, such as intolerance and toxicity. For these reasons, it is crucial to find new drugs to treat PCP.

Echinocandins constitute an alternative class of antifungal drugs to consider for the treatment of PCP. This class includes caspofungin (CAS), anidulafungin, and micafungin. They are cyclic hexapeptides with fatty acyl side chains and act as non-competitive inhibitors of the catalytic subunit Gsc1 of the 1,3-β glucan synthase enzymatic complex (10). β-glucan molecules are components of the cell wall that are homopolymers of β-1,3 linked D-glucose with β −1,6 linked D-glucose side chains present in minority. The Gsc1 protein of *Pneumocystis carinii*, the species infecting rats, was reported to be inhibited by the compound L-733,560, a molecule structurally close to echinocandins (11). Recently, we identified and functionally ascertained the function of the Gsc1 subunit of *P. jirovecii* using complementation of the orthologous gene of *Saccharomyces cerevisiae* (12). The presence of a unique *gsc1* gene in the genome of *P. jirovecii*, as in that of *P. carinii*, further suggests that the Gsc1 subunit is a potential new drug target to fight PCP.

In *S. cerevisiae*, the 1,3-β glucan synthase catalytic subunit is encoded by two different genes, *GSC1* and *GSC2*. A third paralog, *GSC3*, is also present but it is involved only during sporulation. The two subunits *GSC1* and *GSC2* are functionally redundant, but their expression is differentially regulated. The expression of *GSC1* is constitutive and responsible for the cell wall synthesis during the vegetative growth, while that of *GSC2* is induced by glucose deprivation or pheromones and is also involved in cell wall synthesis during sporulation. *GSC1* and *GSC2* genes have an essential overlapping function, *i.e*., only disruption of both genes is lethal. Importantly, the *GSC2* gene can replace the function of the *GSC1* gene during vegetative growth in the case of loss by mutation or deletion (13). The *S. cerevisiae* strain with a deletion of the *GSC1* gene shows a reduced and impaired growth in presence of CAS (14) or anidulafungin (15), but not of micafungin (15). On the other hand, the *S. cerevisiae* wild type shows a normal growth in presence of low doses of CAS and anidulafungin, but its growth is severely impaired in presence of micafungin. These observations showed that the *S. cerevisiae* Gsc1 and Gsc2 subunits have different sensitivities to each echinocandin despite that their identity at the amino acids sequence level is as high as 87% over the whole protein, with 81 and 94% identity at the level of the 1,3-β glucan synthase domains 1 and 2, respectively (Table 1). To our knowledge, the polymorphisms responsible for these different sensibilities have not been determined so far.

**TABLE 1.** Sequence identity (%) of Gsc proteins to their orthologs and paralogs

| | | Whole protein | 1,3-β glucan synthase domain 1 | 1,3-β glucan synthase domain 2 |
| --- | --- | --- | --- | --- |
| *P. jirovecii* Gsc1<sup>a</sup> to | *P. carinii* Gsc1<sup>a</sup> | 90 | 94 | 97 |
|  | *P. murina* Gsc1<sup>a</sup> | 91 | 95 | 96 |
|  | *S. cerevisiae* Gsc1<sup>a</sup> | 59 | 70 | 73 |
| *S. cerevisiae* Gsc1<sup>b</sup> to | *S. cerevisiae* Gsc2<sup>b</sup> | 87 | 81 | 94 |
|  | *S. cerevisiae* Gsc3<sup>b</sup> | 51 | 57 | 60 |

<sup>a</sup> Alignment of these proteins is shown in Figure S1.
<sup>b</sup> Alignment of these proteins is shown in Figure S2.

Spontaneous mutants resistant to echinocandins were initially isolated in *S. cerevisiae* and *Candida albicans* (16–18). Rare clinical isolates of *C. albicans* were also found to be resistant (19, 20). A specific substitution of a serine in position 645 to a proline (S645P) was identified in all spontaneous and most clinical resistant *C. albicans* isolates (20). It is localized within a highly conserved region of the Gsc1 protein in which other mutations conferring resistance to CAS were also identified in *C. albicans* (20). This “Hot Spot no. 1” of mutations starts at residue 641 and ends at residue 649 of *C. albicans* Gsc1. A second but less relevant hotspot of mutations conferring resistance has been identified in another region of the enzyme, from residue 1357 to residue 1364. The S645P substitution has been most frequently observed, F641S being the second most frequent substitution (21). The mutation corresponding to the *C. albicans* S645P substitution introduced by site-directed mutagenesis was found to confer reduced susceptibility to CAS *in vitro* to the mould *Aspergillus fumigatus* (22, 23).

Although demonstrated against *P. carinii* and *P. murina* in rodent models (10, 24–28), the efficacy of CAS against *P. jirovecii* remains controversial. Indeed, clinical reports documented the clearance of PCP treated with CAS alone (29–31), or used in combination with cotrimoxazole (32–36) or clindamycin (37). However, failures of CAS treatment were also described (38, 39). Despite the generally high conservation of active sites among orthologous enzymes, one cannot exclude that the sensitivity to CAS may vary among *P. jirovecii* and the two *Pneumocystis* species infecting rodents because these species are relatively distant from each other (20% of mean divergence at nucleotide level in genomic coding sequences, 40). The *P. jirovecii* Gsc1 subunit bears 90 and 91% identity with those of *P. carinii* and *P. murina*, respectively (Table 1). At the level of the 1,3-β glucan synthase domains 1 and 2, *i.e*., the active sites, the identities are from 94 to 97%. These values are comparable to those between the Gsc1 and Gsc2 subunits of *S. cerevisiae* (see above), which present drastically different sensitivities to the different echinocandins.

The aim of the present study was to determine if the Gsc1 subunit of *P. jirovecii* is sensitive to the echinocandin CAS, as those of *P. carinii* and *P. murina*. To investigate the issue, we analysed the level of sensitivity of *S. cerevisiae* strains functionally complemented by the expression of the wild-type or mutated enzymes of the three *Pneumocystis* species.

## MATERIALS AND METHODS

### Strains and growth conditions

Y05251 is a *S. cerevisiae* haploid strain in which the 1,3-β glucan synthase catalytic subunit gene *GSC1* (also called *FKS1*) was deleted (*MATa his3Δ0 leu2Δ0 met15Δ0 urα3Δ0 YLR342w::kanMX4*). It was obtained from Euroscarf (European *S. cerevisiae* Archive for Functional Analysis [http://www.euroscarf.de]). The strain, named the *gsc1* deletant hereafter, exhibits a slow growth rate and an impaired growth in the presence of low doses of CAS (14). The parental strain of the *gsc1* deletant is BY4741 (*MATa his3Δ1 leu2Δ0 met15Δ0 urαΔO*), and was also obtained from Euroscarf (hereafter named wild-type, WT). The latter was used as a control in the sensitivity tests and in MIC assays. Strains were grown on complete yeast extract-peptone-dextrose (YEPD) medium (1% wt/vol Difco yeast extract, 2% DIfco peptone, 2% glucose).

*Candida albicans* (ATCC 10231) and *Candida parapsilosis* (ATCC 22019) were purified on Sabouraud medium (0.5% wt/vol casein peptone, 0.5% meat extract peptone, 2% glucose), and then grown on minimal solid yeast nitrogen base (YNB) medium (0.67% wt/vol yeast nitrogen base, 2% glucose, 2% Gibco agar) supplemented with a complete supplement mixture (CSM, MP Biomedicals).

### Cloning of the fungal *gsc1* genes

To identify the *P. murina gsc1* gene, the *P. carinii* Gsc1 protein (ID Q9HEZ4) was used as query sequence in BLASTp search against *P. murina* proteome at http://blast.ncbi.nlm.nih.gov/Blast.cgi. A single putative ortholog was detected (locus tag PNEG_03180). The *P. murina* gene sequence encoding the Gsc1 protein was then retrieved from the European Nucleotide Archive (http://www.ebi.ac.uk/ena). The cloning of the *P. jirovecii* and *S. cerevisiae gsc1* genes was previously described (12). Since the *P. carinii* and *P. murina gsc1* genes include each three introns, their cDNAs were synthesized and cloned into the p416GPD vector (41) by GeneCust Europe (Ellange Luxemburg). Their size without introns is respectively 5835 bps and 5847 bps.

To perform a control of sensitivity of our heterologous expression model, the *GSC1* genes of *C. albicans* (GenBank D88815) and *C. parapsilosis* (EU221325) were amplified by PCR from yeast genomic DNA extracted as described previously (42). The detailed procedures of PCR amplification using the proofreading high-fidelity Expand polymerase (Roche Diagnostics) and cloning were described previously (43). Their sizes are respectively 5694 and 5730 pbs. PCR primers and conditions are listed in Tables S1 and S2. Because these primers were intended for oriented cloning, they were designed to create unique restriction sites at ends of the PCR products. After the PCR reactions, the products were extracted using the QIAquick gel extraction kit (Qiagen, Basel, Switzerland). For cloning each *Candida GSC1* gene into the p416GPD expression vector, the double restriction described in Table S1 were used.

### Site-directed mutagenesis

The Gsc1 protein sequences of *C. albicans* (UniProt ID O13428), *S. cerevisiae* (P38631), *P. jirovecii* (L0PD34, locus tag PNEJI1_001061), *P. carinii* (Q9HEZ4), *P. murina* (M7P3D9, locus tag PNEG_03180), and *C. parapsilosis* (A9YLC3) were aligned using T-Coffee (44). This alignment allowed determining the positions within the *Pneumocystis* genes corresponding to the mutations F641S and S645P conferring resistance to CAS in *C. albicans* (Fig. 2; alignment of the complete proteins is shown in Fig. S1). To perform site-directed mutagenesis, two different kits were used. The QuikChange II XL Site-Directed Mutagenesis Kit (Agilent Technologies) was used to create the mutation in the *P. jirovecii gsc1* gene leading to the substitution of the serine at position 718 of the Gsc1 protein into a proline (S718P). The Q5 Site-Directed Mutagenesis Kit (BioLabs) was used to introduce the F714S/S718P double substitution in *P. jirovecii*, the S715P substitution in *P. carinii*, and the S719P substitution in *P. murina*. Mutagenesis were performed according to manufacturers’ instructions. Minipreparations of plasmid DNA were subsequently carried out (45). In order to verify the presence of the desired mutations, an internal segment of the *gsc1* genes was amplified and subsequently sequenced. Primers for mutagenic reactions and PCR amplifications are listed in Table S1. Mutagenesis amplification reactions and PCR conditions are described in Table S2. Sequencing of both strands was performed thanks to the two primers used for amplification, as well as the BigDye Terminator DNA sequencing kit and ABI PRISM 3100 automated sequencer (both from PerkinElmer Biosystems).

### Transformation of the *S. cerevisiae gsc1* deletant

Transformation with plasmids containing the *P. jirovecii gsc1* or the *S. cerevisiae GSC1* gene were previously described (12). The *S. cerevisiae GSC1* gene could not be cloned in the p416GPD plasmid because of restriction sites issues, but into p415GPD (leu marker instead of ura). The recombinant p416GPD plasmids containing the *Pneumocystis* mutated *gsc1* alleles, as well as the *C. albicans* or *C. parapsilosis GSC1* gene were introduced into the *gsc1* deletant by transformation for uracil prototrophy using the one-step method (46). Transformants were selected on solid YNB medium supplemented with CSM (MP Biomedicals) lacking uracil. In order to be used as controls in the sensitivity tests and in the MIC assays, the *gsc1* deletant and the WT were transformed with the empty p416GPD plasmid. Three transformants clones of each constructed strain were randomly chosen and purified by growth on the same selective medium.

### Test of complementation and susceptibility to caspofungin

Before studying the sensitivity to echinocandins, we had to assess the function of the *P. carinii* and the *P. murina gsc1* genes, as we previously carried out for the *P. jirovecii gsc1* gene (12). Functional complementation of the *gsc1* deletant was proven by the spot dilution test on YNB selective medium lacking uracil and supplemented with or without 150 ng/ml CAS (Fluka, Chemie AG). To that aim, transformant isolates carrying the *P. carinii gsc1* or *P. murina gsc1* gene were grown overnight in selective medium YNB supplemented with CSM lacking uracil to avoid the loss of the plasmid. Cells were then diluted at an optical density (OD) at 540 nm of 0.1 in NaCl 0.9% wt/vol (ca. 7.5 x 10^5^ cells/ml). Four serial 10-fold dilutions in NaCl 0.9% were prepared, and 3 μl of each dilution were spotted on the medium. Spots were observed after 3 to 4 days of incubation at 30°C. The same procedure assessed the functionality and sensitivity to CAS of the strains complemented with the mutated *gsc1* alleles.

### MIC assessment using E-test^®^

Each strain was grown overnight in selective medium YNB + CSM lacking uracil, or leucine for *S. cerevisiae GSC1* gene, and then adjusted in NaCl 0.9 % to an OD540=0.2 (~1.5 x 10^6^ cells/ml). One hundred microliters of this dilution was spread on fresh YNB solid medium + CSM lacking uracil or leucine. A single strip of E-test caspofungin (Biomérieux) was then applied on each petri dish. MICs were read after 2 days of incubation at 30°C, or 35°C for the *Candida* species. The MIC was defined as the concentration at which no growth was observed on both sides of the E-test strip.

### MIC assessment using Sensititre YeastOne^®^

Each strain was grown overnight in selective medium YNB, then adjusted in NaCl 0.9 % to an OD540=0.2 (ca. 1.5 x 10^6^ cells/ml). Twenty microliters of this dilution were then diluted into 11 ml of YeastOne^®^ inoculum broth in order to obtain ca. 3 x 10^3^ cells/ml. One hundred microliters was then transferred into each well of a YeastOne^®^ plate (Thermofisher Scientific). Plates were observed and MICs determined after 36 h of incubation at 30°C, or 35°C for the *Candida* species. MIC was defined as the first well in which no pellet of cells was observable.

## RESULTS

Functional ascertainment of the *P. carinii* and *P. murina gsc1* genes by complementation of the *S. cerevisiae gsc1* deletant. As previously reported for *P. carinii* (11) and *P. jirovecii* (12), we identified a single Gsc1 protein within the *P. murina* proteome by a homology search using the Gsc1 protein of *P. carinii* as the query sequence. To ascertain the function of the *P. carinii* and *P. murina gsc1* genes, recombinant plasmids expressing them were introduced into the *S. cerevisiae gsc1* deletant. As controls, an empty plasmid was introduced in the WT and *gsc1* deletant strains, and a plasmid expressing the *S. cerevisiae GSC1* gene was introduced in the deletant. The identities of the Gsc1 proteins studied relatively to that of *S. cerevisiae* are given in Table S3. Serial dilution of the transformed strains were spotted on medium containing or not CAS (spot dilution test, Fig. 1). The deletion of the *GSC1* gene in *S. cerevisiae* causes a paradoxical effect, i.e., an increased susceptibility to CAS, though the target of CAS is absent (14). This is due to the replacement of Gsc1 by Gsc2, an enzyme that is more sensitive to CAS (13). On medium supplemented with CAS, a complete restoration of the wild-type growth was observed in presence of the control *S. cerevisiae GSC1* gene, but not in the presence of the empty vector (Fig. 1). A partial restoration was observed in the presence of the *P. carinii* or *P. murina* gene, as we previously reported for *P. jirovecii* and reproduced here (Fig. 1). These observations demonstrated that the expression of *P. carinii* and *P. murina gsc1* genes rescued the function of the deleted *S. cerevisiae GSC1* gene, ascertaining their function. In order to investigate the sensitivity to CAS of the three *Pneumocystis* enzymes, we used site-directed mutagenesis to introduce mutations that correspond to those conferring resistance in other fungi.

**FIG 1.**
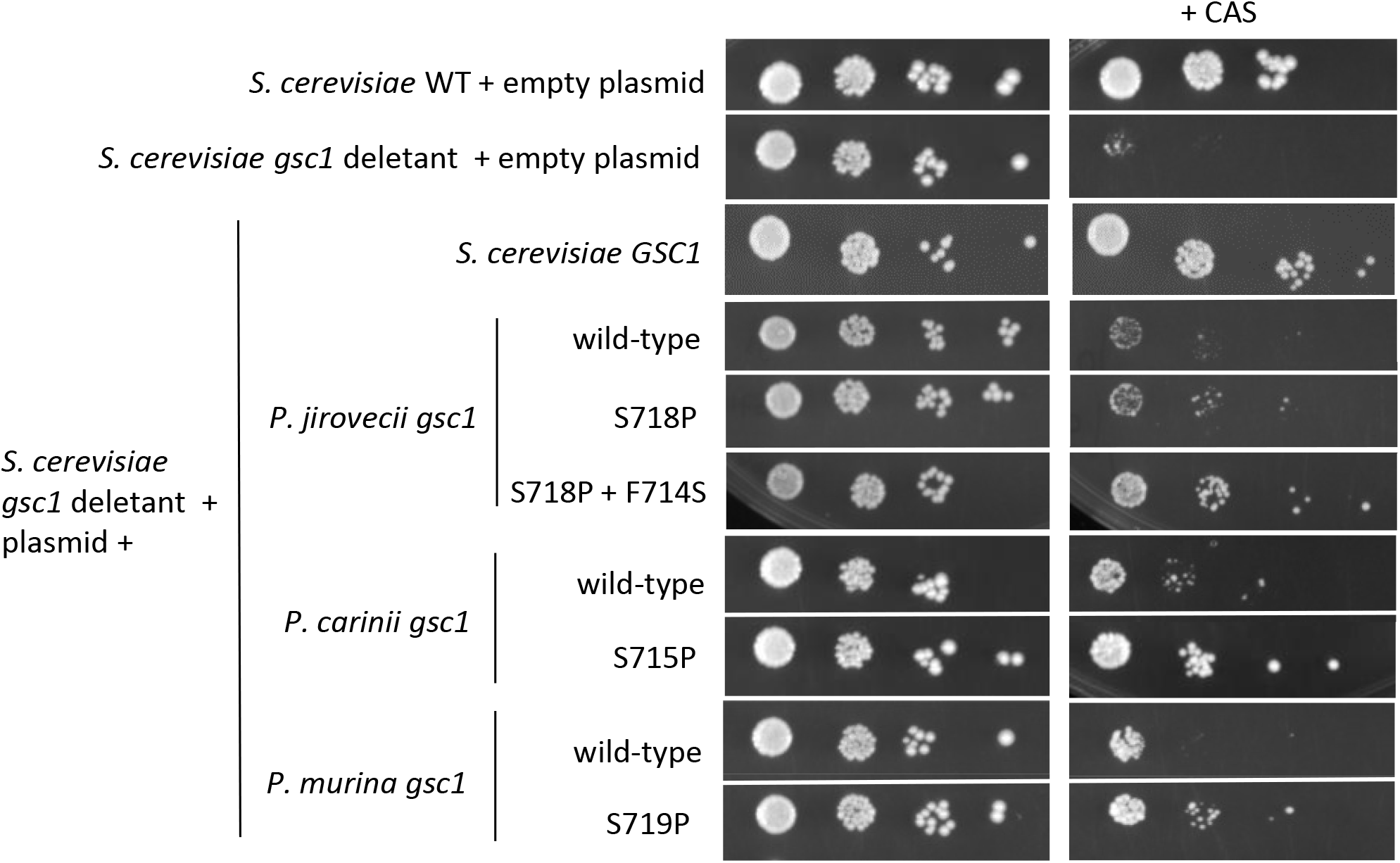
Sensitivity to caspofungin (CAS) of *S. cerevisiae* WT and functionally complemented *gsc1* deletant strains using the spot dilution test. The complementing genes expressed on plasmid, wild-type or encoding the indicated amino acid substitution, are described on the left. Log dilution of a suspension of cells at ca. 7.5 x10^5^/ml were spotted on minimal selective medium without (left) or with (right) 150 ng/ml of CAS, and incubated for 3 days at 30°C. The most concentrated suspension is on the left. The complementing gene was expressed on plasmid p416GPD except that of *S. cerevisiae* on p415GPD because of restriction sites issues. Minimal selective medium YNB supplemented with CSM without uracil was used to select for p416GPD, while YNB supplemented with CSM without leucin was used to select p415GPD. Three independent isolates of each strain were analyzed, one representative isolate is shown here.

### Sensitivity to CAS of the *S. cerevisiae* strains complemented with the *Pneumocystis* Gsc1 mutated proteins

Mutants resistant to echinocandins carrying mutations F641S and S645P within the hotspot no. 1 of Gsc1 have been described in the pathogenic fungus *C. albicans* (16–20). The sequences of this hotspot of mutations of the *P. jirovecii, P. carinii*, and *P. murina* Gsc1 protein were aligned with those of *C. albicans, S. cerevisiae*, and *C. parapsilosis* (Fig. 2; alignment of the complete proteins is shown in Fig. S1). This alignment identified the positions in the three *Pneumocystis gsc1* genes corresponding to the *C. albicans* F641S and S645P substitutions. Site-directed mutagenesis was used to introduce one or two mutations encoding the corresponding substitutions within the *gsc1* gene of *P. jirovecii, P. carinii*, or *P. murina* (the polymorphisms introduced at the nucleotide sequence level are described in Table S1).

**FIG 2.**
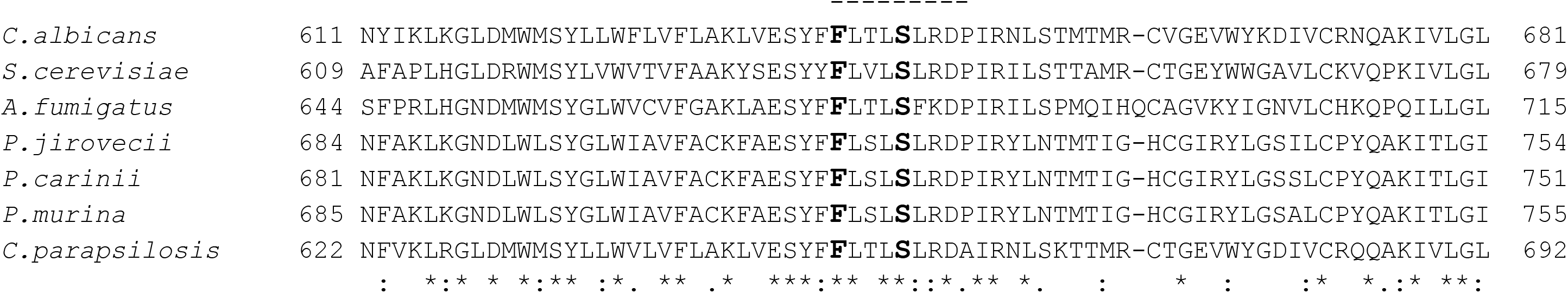
Multiple-sequence alignment of the region encompassing the hot spot no. 1 of mutations of Gsc1 proteins. T-Coffee was used (44). The identical, strongly, and weakly conserved residues are indicated by asterisks, double points, and single points, respectively. Dashes indicate gaps. The hot spot no. 1 of mutations (20) is shown above the alignment by the dashed line. Residues F641 and S645 conferring CAS resistance in *C. albicans* and the corresponding residues in the other proteins are in bold. The natural resistance of *C. parapsilosis* is due to the polymorphism P660A at the end of the same hotspot of mutations. The alignment of the complete proteins is show in Figure S1.

The partial restoration of the wild-type growth on CAS observed with the *P. jirovecii* Gsc1 enzyme increased in presence of one mutation, and increased more upon introduction of the two mutations simultaneously (Fig. 1). Similarly, the partial restoration with the *P. carinii* or *P. murina* enzyme increased in presence of a single mutation. This increase of complementation efficiency corresponds to a decrease of sensitivity to CAS. This demonstrated that the three wild-type *Pneumocystis* enzymes present a certain level of sensitivity to CAS.

### Minimum inhibitory concentration (MIC) assessment using E-test^®^ and Sensititre YeastOne^®^

We determined the MICs of CAS for the *S. cerevisiae* WT and complemented *gsc1* deletant strains. To assess the sensitivity of the two methods, we also analysed *C. albicans* and *C. parapsilosis*, as well as the *S. cerevisiae gsc1* deletant complemented with the *GSC1* gene of these two *Candida* species. The former *Candida* species is susceptible to CAS, whereas the latter is considered as resistant (47, 48). The natural resistance of *C. parapsilosis* is due to a polymorphism at the end of the hotspot no. 1 (P660A), which has not been observed in *C. albicans* so far (49; Fig. 2). In agreement with the spot dilution tests described here above, we observed using both E-test^®^ and YeastOne^®^ an increased susceptibility of the *S. cerevisiae gsc1* deletant compared to the WT (respectively 0.125 and 0.12 versus 0.250 and 0.25 μg/ml; Table 2; the E-test results are shown in Fig. S3). All *S. cerevisiae* strains complemented with the *Pneumocystis* wild-type or mutated genes had MICs identical to those of the *gsc1* deletant (0.125 for E-tests and 0.12 for Yeastone). The decreased sensitivity to CAS conferred by the mutations introduced was not detected using both methods. Thus, these methods are less sensitive than the spot dilution test used above since the latter always allowed detection of this decrease in several experiments. The MIC values for *C. albicans* whole cells using E-test^®^ and YeastOne^®^ were similar to those for the *S. cerevisiae* WT strain (0.380 and 0.12 versus 0.250 and 0.25), whereas, consistently with its resistance to CAS, *C. parapsilosis* had higher MICs using both methods (0.500 and 8.00). The decreased sensitivity of *C. parapsilosis* was also detected using both methods upon heterologous expression of its Gsc1 subunit in *S. cerevisiae* (0.190 and 0.25 versus 0.125 and 0.12 for *C. albicans* Gsc1). Using the heterologous expression system, the wild-type *Pneumocystis* Gsc1 subunits resulted in identical MICs than the *C. albicans* Gsc1 (0.125 and 0.12), whereas *C. parapsilosis* Gsc1 presented higher MICs (0.190 and 0.25). These observations suggested that the sensitivity level to CAS of the three *Pneumocystis* enzymes is similar to that of *C. albicans*, which is susceptible to CAS.

**TABLE 2.** Minimum inhibitory concentrations (MIC) of caspofungin (CAS) for the *S. cerevisiae* WT and functionally complemented *gsc1* deletant strains, as well as for *Candida* species^a^

| Strain | MIC (µg/ml) |  |
| --- | --- | --- |
|  | E-test | YeastOne |
| *S. cerevisiae* WT + empty plasmid | 0.250 | 0.25 |
| *S. cerevisiae gsc1* deletant + empty plasmid | 0.125 | 0.12 |
| *S. cerevisiae gsc1* deletant + plasmid + *S. cerevisiae GSC1* | 0.250 | 0.25 |
| *P. jirovecii gsc1* wild-type | 0.125 | 0.12 |
| S718P | 0.125 | 0.12 |
| S718P + F714S | 0.125 | 0.12 |
| *P. carinii gsc1* wild-type | 0.125 | 0.12 |
| S715P | 0.125 | 0.12 |
| *P. murina gsc1* wild-type | 0.125 | 0.12 |
| S719P | 0.125 | 0.12 |
| *C. albicans GSC1* | 0.125 | 0.12 |
| *C. parapsilosis GSC1* | 0.190 | 0.25 |
| *C. albicans* | 0.380 | 0.12 |
| *C. parapsilosis* | 0.500 | 8.00 |

<sup>a</sup> One isolate among three of each complemented strain was chosen randomly for analysis. One out of two experiments that gave similar results is reported here. Although obtained using various methods, the MIC values previously published correspond roughly to ours for *S. cerevisiae* WT (0.25 versus 0.03-0.4; 14, 15, 20) and *gsc1* deletant (0.12-0.125 versus 0.0015-0.1; 14, 15, 20), as well as for *C. albicans* (0.12-0.380 versus 0.12-0.25; 20, 47) and *C. parapsilosis* (0.500-8.00 versus 0.25-8; 47-49).

## DISCUSSION

Because of the absence of an *in vitro* culture method, sensitivity to CAS cannot be performed on whole *Pneumocystis* cells. Consequently, we studied the Gsc1 enzymes of three *Pneumocystis* species in the heterologous system of expression of *S. cerevisiae*. We used site-directed mutagenesis to introduce into the *Pneumocystis* enzymes the substitutions corresponding to those conferring resistance to CAS in *C. albicans*. This revealed that, despite the divergence among their active sites, the three *Pneumocystis* Gsc1 enzymes are susceptible to CAS, and this to a similar level. Because CAS has been demonstrated to be effective against *P. carinii* and *P. murina* in rodent models (26, 27), this observation suggested that CAS is also effective against *P. jirovecii*. Moreover, MICs determination showed that the level of sensitivity of *Pneumocystis* Gsc1 was similar to that of the *C. albicans* enzyme, suggesting that the sensitivity of the *Pneumocystis* enzymes is at a level that is usable clinically. It is of course difficult to translate our results obtained at the enzyme level to the whole cell level. Nevertheless, Gsc1 is a cell surface enzyme that is easily reachable by drugs, and thus more likely to behave similarly among the three *Pneumocystis* species. A structural difference of the cell wall could induce varying sensitivity to CAS of the Gsc1 subunit among the three *Pneumocystis* species. However, there is presently no obvious reasons to think that the wall of *P. jirovecii* is different from those of *P. carinii* and *P. murina*. Finally, our results support the high relevance of the animal models as tools to understand the effect of CAS on the human pathogen *P. jirovecii*.

Studies in animal models showed that echinocandins provoke the disappearance of *P. carinii* and *P. murina* asci but not of the trophic forms, probably because the latter cells have no or little cell wall made of 1,3-β glucan (26). Thus, the treatment did not eradicate the infection, and its cessation resulted in the repopulation in asci from the remaining trophic cells. Consequently, it is likely that CAS is useful only if used in combination with another therapy targeting trophic forms, or both cellular forms such as cotrimoxazole. CAS inhibited efficiently the dissemination of the pathogen in animal models (26), which is consistent with the fact that asci are believed to be the transmission particles (26, 50). This feature of CAS might prove useful in the management of infected and susceptible patients within the hospital by inhibiting interhuman transmission of *P. jirovecii*.

In conclusion, our results demonstrate that the human pathogen *P. jirovecii* Gsc1 enzyme is sensitive to caspofungin, similarly to the enzymes of the animal pathogens *P. carinii* and *P. murina*. This suggests that echinocandins might be a good alternative to treat PCP in humans when used in combination with an established treatment. The use of echinocandins to fight *Pneumocystis* infections deserves further investigation.

## ACKNOWLEDGMENTS

This work was supported by Swiss National Science Foundation grant 310030_165825. The present work was submitted by A. Luraschi as a partial fulfilment for a PhD degree at the Faculty of Biology and Medicine of the University of Lausanne. We thank Michel Monod for his critical reading of the manuscript.

## SUPPLEMENTAL MATERIAL

**Table S1**. Oligonucleotide primers used for mutagenesis reactions and PCR amplifications

**Table S2**. Conditions of mutagenesis and PCR reactions

**Table S3**. Sequence identity (%) of the *S. cerevisiae* Gsc1 protein to the ortholog proteins studied in the complementation assays and MIC determinations

**Fig. S1**. Multiple sequence alignment of Gsc1 proteins of relevant fungi. T-Coffee was used (44). The identical, strongly, and weakly conserved residues are indicated by asterisks, double points, and single points, respectively. Dashes indicate gaps. The hot spots no. 1 (20) and 2 (49) of mutations, as well as the 1,3-β glucan synthase domains 1 and 2, are indicated above the alignment. Residues F641 and S645 conferring CAS resistance in *C. albicans* and the corresponding residues in the other proteins are shown in bold. The natural resistance of *C. parapsilosis* is due to a polymorphism at the end of the hotspot no. 1 (P660A).

**Fig. S2**. Multiple sequence alignment of *S. cerevisiae* Gsc proteins. The Uniprot accession numbers of Gsc1, Gsc2, and Gsc3 are respectively P38631, P40989, and Q04952. T-Coffee was used (44). The identical, strongly, and weakly conserved residues are indicated by asterisks, double points, and single points, respectively. Dashes indicate gaps. The 1,3-β glucan synthase domains 1 and 2 are indicated above the alignment.

**Fig S3**. E-test determination of the minimum inhibitory concentration (MIC) of caspofungin (CAS) for the *S. cerevisiae* WT and functionally complemented *gsc1* deletant strains, as well as for *Candida* species. One hundred microliters of a suspension of cells at ca. 1.5 x 10^6^ cells/ml were spread on minimal selective medium lacking uracil, or lacking leucin for the *S. cerevisiae GSC1* gene. The CAS E-test strip was deposited, and the plate was incubated at 30°C, or 35°C for the *Candida* species. The concentration at which no growth was observed on both sides of the E-test strip was defined as the MIC.

